# Coacervates protect RNA against hydrolysis under wet-dry cycling conditions

**DOI:** 10.64898/2026.08.03.742545

**Authors:** Jiaqi Pei, Philip C. Bevilacqua, Christine D. Keating

## Abstract

RNA is crucial to all extant biology and thought to have played an essential role in life’s origins. However, hydrolysis of RNA under common environmental conditions such as heating, drying, and exposure to divalent cations presents challenges for its persistence and effectiveness in prebiotic contexts. RNAs with less stable secondary and tertiary structures are especially susceptible to degradation. In living cells, sequestration in biopolymer-rich membraneless organelles helps shield RNA from damage and enzymatic degradation. Here, we report that compartmentalization into polyelectrolyte-based complex coacervate droplets protects RNA from hydrolysis during wet-dry cycling at elevated temperatures in the presence of Mg^2+^. This protection can be understood in terms of the coacervate microenvironment under varied salt concentrations and how it changes during sample drying. After gaining insight into this prebiotically-plausible mechanism using an unstructured model RNA, we demonstrate coacervate-based protection of a ligase ribozyme, supporting the functional relevance of the protection.

## Introduction

RNA is necessary for all extant life and thought to have been critical to life’s emergence. Its plausible abiotic production, ability to not only store information but also perform catalysis, and central role in translation via the ribosome support the importance of RNA prior to the origin of life^1–4^. However, RNA can be quite susceptible to chemical damage such as hydrolytic cleavage of its backbone phosphodiester bonds, particularly outside the protective environment of living cells^5, 6^. RNA degradation is favored by conditions probable on the early Earth: (1) Many early RNAs likely lacked stable folding, making them more susceptible to cleavage (2) Mg^2+^ was likely available in prebiotic waters. Although Mg^2+^ can be beneficial for RNA folding and ribozyme activities, it catalyzes RNA hydrolysis (3) Elevated temperatures, frequently invoked as part of wet-dry cycling to drive oligomerization reactions in prebiotic scenarios, also increase hydrolysis rates^7^. Considering these challenges, it is critical to study whether there are potential prebiotic mechanisms that could stabilize RNAs against hydrolysis. We chose to investigate the effect of wet-dry cycling on hydrolysis of an unstructured RNA in the presence of Mg^2+^.

Wet-dry cycling is a prebiotically relevant geological process of repeated heating and evaporation, followed by rehydration that can promote molecular complexity^8–11^, including RNA synthesis^9, 10, 12^. However, RNA oligomers produced by wet-dry cycling are, in general, considerably shorter (∼10mer) than functional ribozymes (> 50mer)^12–16^. Indeed, RNA hydrolysis limits chain growth progress made by condensation during drying^15, 17^. This leaves a question: How could sufficient prebiotic RNA have escaped hydrolysis and grown long enough to adopt functional three-dimensional structures? Here, we propose that compartmentalization could provide an answer.

Formation of membraneless organelles by associative liquid-liquid phase separation, or coacervation, is a common means of RNA compartmentalization in extant biology. Indeed, structures such as stress granules and aggresomes preserve intracellular RNA during stress^18, 19^, and phase separation has been proposed as contributing to desiccation tolerance in anhydrobiotes such as seeds or tardigrades^20, 21^. Coacervate droplets are also plausibly prebiotic compartments due to their ease of formation from a wide range of molecular components (including peptides, nucleic acids, carbohydrates as well as nonbiological polymers)^22–25^ and ability to recruit and concentrate prebiotically important cargo, such as Mg^2+^ and RNA ^26–28^. These compartments serve as alternative microenvironments with properties dependent on their molecular composition and tunable over a wide range of physicochemical and functional properties^29, 30^.

Notably, increased ribozyme reaction rates and nonenzymatic RNA extension have been reported in coacervates^3, 31–39^. Previous work showed how changes in coacervate physical and chemical properties during the drying process could be understood on the basis of the phase diagram^24^. We wondered how RNA guest molecules in such a system would experience wet-dry cycling, and whether the coacervate microenvironment might provide protection from degradation, thus contributing to the origin of life.

Here, we report that RNA hydrolysis at elevated temperature during wet-dry cycling is reduced upon encapsulation within coacervates. We elucidate how the coacervate microenvironment provides this protection by means of its molecular composition, physical properties, interaction network, and chelation of Mg^2+^, and how polymer and salt concentrations influence the extent of RNA hydrolysis. Experimental RNA fragmentation profiles after multiple wet-dry cycles can be understood in terms of reduced per-bond hydrolysis probability in the presence of coacervates. Finally, we show that a ligase ribozyme retains functional integrity after wet-dry cycling in the presence of coacervates, overcoming loss of activity in their absence. These findings support the potential importance of coacervate-based protocells during environmental wet-dry cycling prior to the emergence of life.

## Results and Discussion

### Coacervates sequester polyelectrolytes and RNA

As our coacervate system to examine RNA stability, we chose a well-characterized polyelectrolyte pair with relatively short polymer lengths – the polycationic poly(diallyldimethylammonium chloride) (PDADMAC, 8.3k, ∼53mers) and the polyanionic polyacrylic acid (PAA, 1.8k, ∼ 25mers) (Fig. 1a-b). Together, these oppositely charged polyelectrolytes undergo complex coacervation, resulting a polymer-rich coacervate phase and a polymer-depleted dilute phase. Samples were prepared with 15 mM in both cationic sidechains from PDADMAC and anionic sidechains from PAA, which we will refer to as 15 mM charge concentration (15 mMc), in 25 mM 4-(2-hydroxyethyl)-1-piperazineethanesulfonic acid (HEPES) buffer (pH 7.5) and 2 mM Mg^2+^. NaCl from polyelectrolyte counterions and pH adjustment are present in these samples (∼20 mM NaCl; see Methods), and we added either 0 mM or 100 mM NaCl to this to generate “low salt” and “high salt” samples for comparison. These mixtures were both visibly turbid. Vibrational spectra collected via MicroRaman show much higher local concentrations of PDADMA and PAA within the droplets as compared to the dilute continuous phase (Fig. 1c-d, Supplementary Table 1). To these coacervate samples, we added 20 nM of polyuracil (polyU 45nt, U45), chosen because it lacks secondary structure and hence all of its phosphodiester bonds should be equally susceptible to hydrolysis^40^. Addition of Alexa488 5’-labeled U45 RNA revealed its accumulation within the coacervates, with confocal fluorescence microscopy showing ∼500-1000-fold higher fluorescence intensity inside the coacervate droplets as compared to the surrounding dilute phase (Fig. 1e, right).

**Figure 1:**
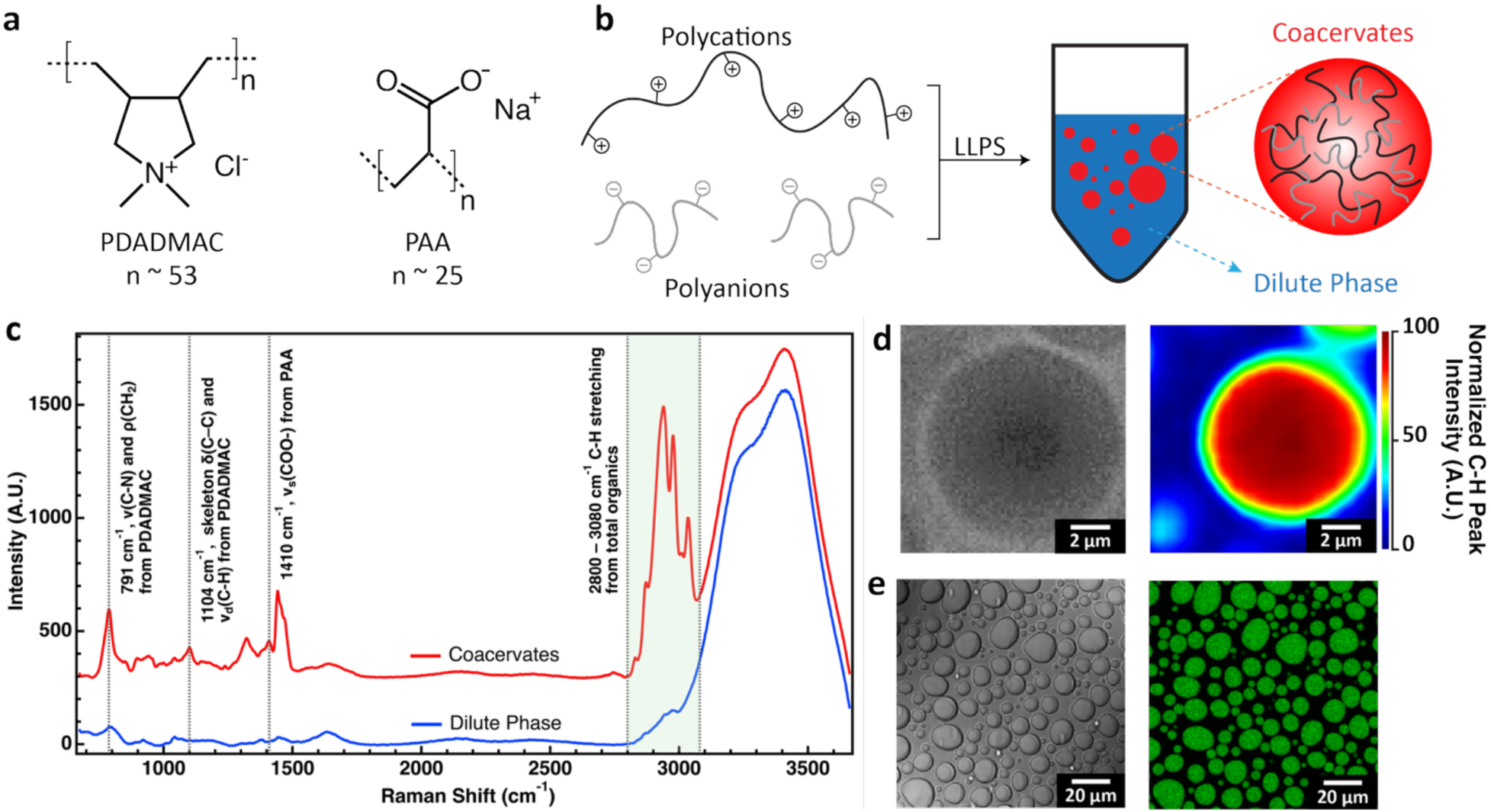
Coacervates sequester polyelectrolytes and RNA. **a** Molecular structures of polymers used to form coacervates. **b** Coacervates were produced by mixing oppositely charged polyelectrolytes. **c** MicroRaman spectra for coacervate phase and dilute phase. Green shaded region represents the C–H stretching (2800 – 3080 cm^-1^) from total organics used in plotting panel d. Coacervate samples in panels **c**-**e** contain 15 mM charge concentration (15 mMc) of PDADMAC and PAA prepared in 25 mM HEPES buffer at pH 7.5, with no added NaCl. Spectra are offset for ease of visualization. **d** MicroRaman image of a coacervate droplet. Left: transmitted light shows droplet location. Right: microRaman map shows accumulation of polyelectrolytes within the coacervate droplet, depicted by normalized intensity for C-H stretching region (2800 – 3080 cm^-1^) of total organics (i.e. PDADMA and PAA). **e** Optical microscope images of PDADMA/PAA coacervate droplets. Left: transmitted light image. Right: fluorescence image showing accumulation of a model RNA (Alexa 488-tagged U45) in the coacervates.

### Phase behavior during dehydration

For each wet-dry cycle, samples were held at 75 °C until dried (180 min for 100 µL samples, Supp. Fig. 1), followed by rehydrating to their original volume (Fig. 2a). The phase diagram (Fig. 2b) helps to explain what happens to the PDADMA/PAA solutions during the drying and rehydration process^24^. Dehydration of an initial composition within the two-phase region (100% volume) can be followed as a diagonal drying trajectory line corresponding to the increasing salt and polyelectrolyte charge concentrations within the sample. Compositions corresponding to our low and high salt samples both cross into the single-phase region prior to reaching complete dryness (Fig. 2b-c). Notably, the initial compositions determine the system’s drying trajectory on the phase diagram. Sample turbidity measured at 75 °C during drying was used to determine when coacervates disappear in both systems (Fig. 2c, Supp. Fig. 2-3). Coacervates in the low salt samples remained until nearly 90% of the initial volume was lost but disappeared after only ∼50% of volume evaporated in the high salt sample. Optical microscopy of these samples, which was measured after cooling to room temperature (Fig. 2c), agreed with the turbidity results.

**Figure 2:**
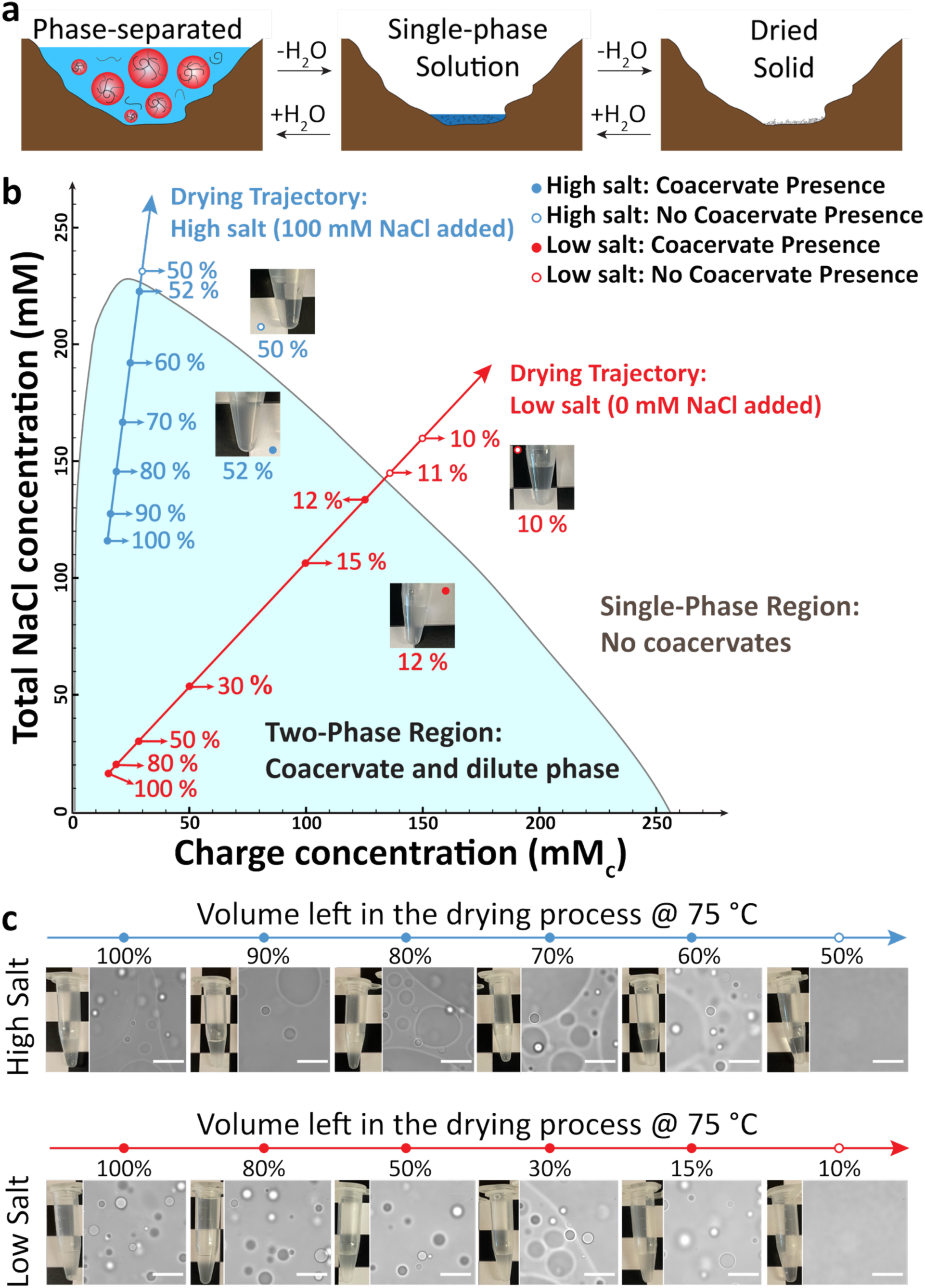
Coacervate system phase behavior during drying. **a** Illustration of wet-dry cycling of coacervate systems in a prebiotic water body. **b** A phase diagram of PDADMA/PAA coacervate system, superimposed on 1:1 charge ratio PDADMA/PAA phase diagram determined at 23 °C by Fares *et al*^24^, is displayed along with experimental drying trajectories. In the phase diagram, the x axis represents polyelectrolyte charge concentration (mM_c_) and y axis represents the total NaCl concentration within the coacervate system. Experimental drying trajectories for the low salt and high salt coacervate systems used here (red and blue lines, respectively). These lines extrapolate to the origin at infinite dilution; only the portion corresponding to our samples’ compositions throughout drying are shown here. Percentage of remaining volume is indicated at points along the drying trajectories, with 100% indicating the starting composition. **c** Photographs (75 °C, left panels) and optical microscopy images (after cooling to room temperature, right panels) for same-volume “mimic” samples with concentrations corresponding to different points along the drying trajectories (see Methods). Scale bars are 20 μm. Additional compositions are shown in Supporting Figures 2-3; these data were used to determine where each sample transitioned from two-phases to single-phase at 75 °C, denoted in (b) as filled and open points, respectively, along the drying trajectories.

**Figure 3.**
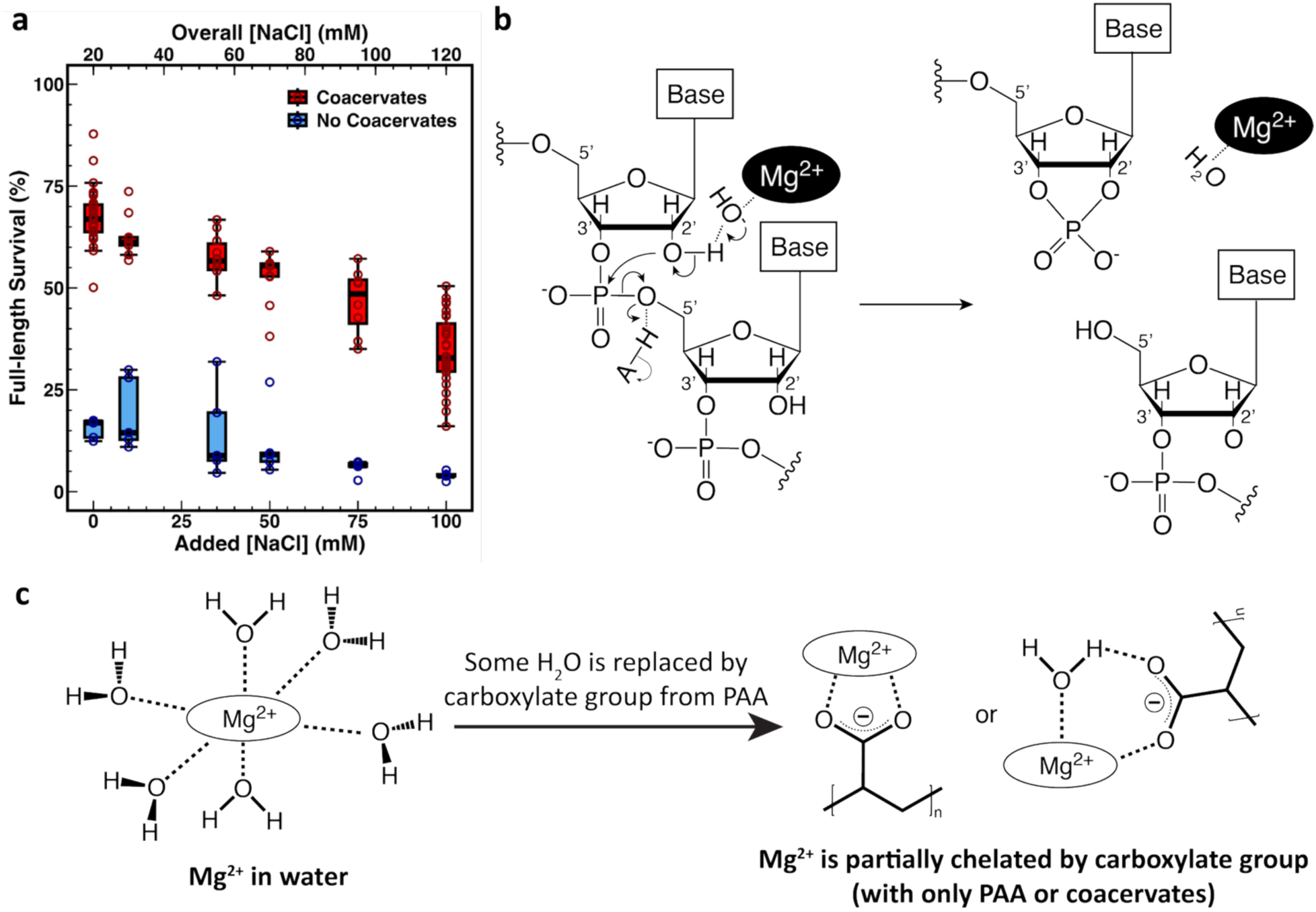
Mg^2+^ catalyzed RNA hydrolysis is suppressed by the presence of coacervates. **a** Full-length survival of U45 after one wet-dry cycle in 2 mM Mg^2+^ with various NaCl concentrations in the presence and absence of coacervates. Samples were held at 75 °C until dried (180 min for 100 µL samples, Supp. Fig. 1), followed by rehydrating to their original volume. **b** Mechanism of Mg^2+^-catalyzed RNA hydrolysis. **c** PAA acts as a weak Mg^2+^ chelator to replace the H_2_O in the hydration shell of Mg^2+^ (not showing the unchanged water after chelation on the left).

Thus, the initial NaCl concentration greatly impacts the extent to which coacervates persist, and hence sequester RNA molecules, during the drying process. Since the loss of coacervates during drying would release previously compartmentalized RNA, we reasoned that these differences in persistence may influence RNA survival.

### Coacervates protect RNA from hydrolysis under wet-dry cycling

To determine how coacervates influence RNA survival, we added 5’-^32^P-labeled U45 and used polyacrylamide gel electrophoresis (PAGE) to visualize the extent of U45 hydrolysis after a single wet-dry cycle (plot in Fig. 3a, gels in Supp. Fig. 4-5). The results showed strong protection of RNA, with coacervate-containing samples retaining >3.5× fold more full-length RNA as compared to no-coacervate samples at low and high salt (Fig. 3a). The highest full-length survival values were found under the lowest salt condition of no added NaCl, with 68 ± 7 % and 15 ± 2% full length remaining in coacervate and no-coacervate samples, respectively. At the highest salt condition of 100 mM NaCl, we found only 35 ± 8% and 4 ± 1% intact U45 RNA after cycling in the coacervate and no-coacervate samples, respectively. These data suggested that coacervates protect sequestered RNA during the wet-dry cycling process at both salt concentrations.

**Figure 4.**
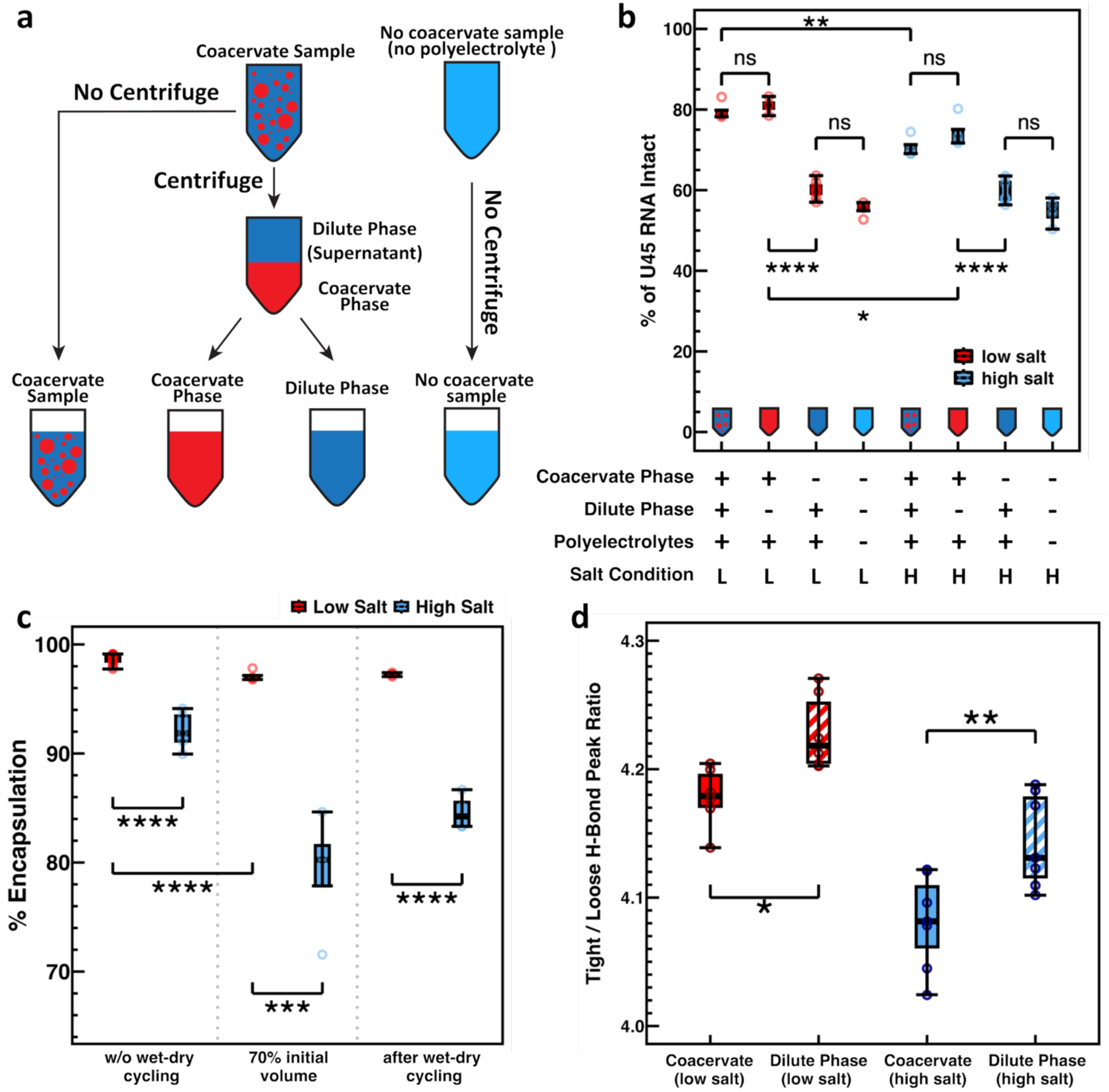
Sequestration within coacervates protects RNA against hydrolysis. **a** Parallel experiments (bottom of flowchart) comparing the amount of RNA hydrolysis among different phases/systems. Experiments were conducted at 75 °C for 180 min under constant volume (2 µL). **b** Full-length survival among different phases/systems illustrated in panel a. In the salt condition row, L represents the low salt condition, and H represents the high salt condition. **c** Encapsulation of U45 within coacervates under different coacervate systems and salt conditions with 5’-^32^P-U45: no wet-dry cycling, same-volume “mimic” samples with concentrations corresponding to point that has 70% initial volume along the drying trajectories, and coacervate system after one wet-dry cycle. **d** The ratio is calculated by the area under 3220 to 3330 cm^-1^(tight H-bond) divided by the area under 3400 to 3420 cm^-1^ (loose H-bond)^50–52^. The data was obtained from a total of 5-6 microRaman spectra across three different repeats. Each spectrum was obtained from a coacervate droplet. t-test statistical significance: ns (no statistical difference, p > 0.05), * (0.01 < p ≤ 0.05), ** (0.001 < p ≤ 0.01), *** (0.0001 < p ≤ 0.001), and **** (p ≤ 0.0001).

**Figure 5.**
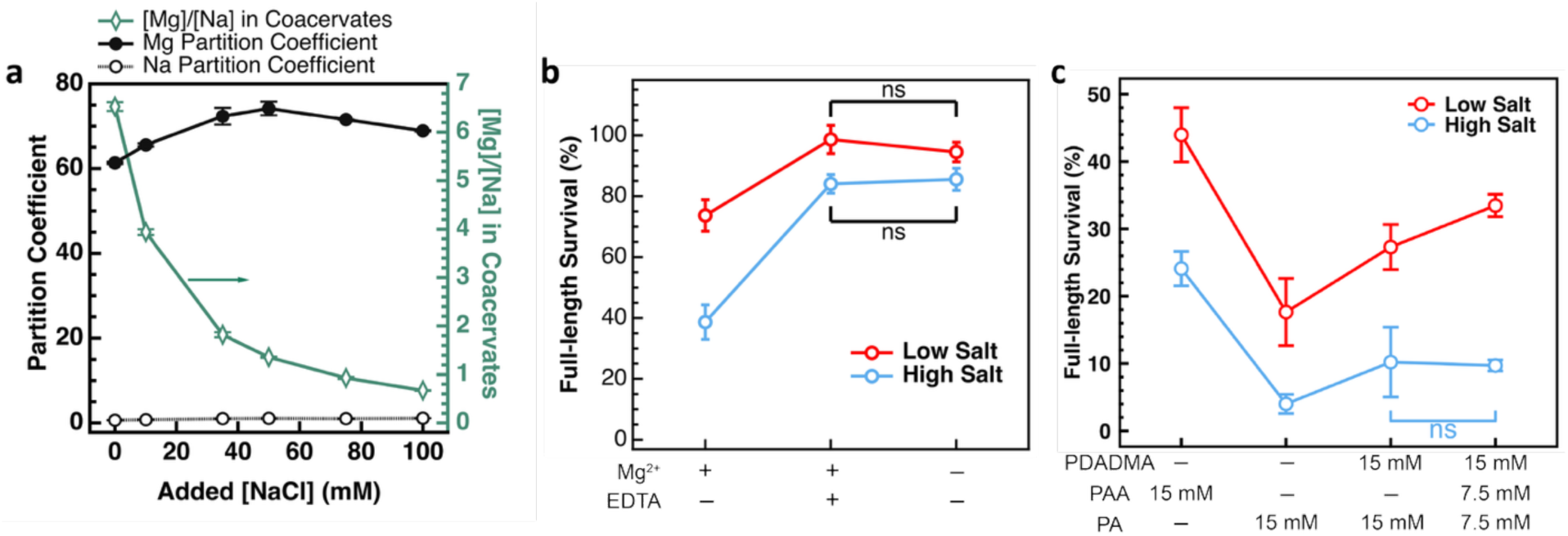
Carboxylate-rich coacervate microenvironment concentrates Mg^2+^ while reducing its ability to hydrolyze RNA. **a** Partition coefficient of Mg^2+^ (solid black line) and Na^+^ (dotted black line) and [Mg^2+^]/[Na^+^] under various added NaCl concentrations, obtained from ICP–AES. +/- represent the item was present/absent in the sample. ns represents two conditions that are not statistical different. **b** Full-length U45 survival in the following systems: coacervates system with added Mg^2+^ (first tick), coacervates system with added Mg^2+^ and EDTA (second tick), and coacervates system with no added Mg^2+^ (third tick). Red and blue represent low and high salt coacervate systems, respectfully. **c** Propionic acid (PA) acted as a weak Mg^2+^ chelator to compare with PAA under the following systems: only 15 mM_c_ PAA and no PDADMAC in the system, only 15 mM_c_ PA and no PDADMAC in the system, 15 mM_c_ PA and PDADMAC in the system, and half of the PA replaced by PAA and PDADMAC in the system (7.5 mM_c_ PAA and 7.5 mM_c_ PAA). The total charge concentration of PA and PAA was kept at 15 mM_c_ under all four conditions.

To understand what role each polyelectrolyte could bring to the observed protection effect, we carried out a series of experiments with only PDADMA or PAA with U45. We found that for a given NaCl concentration, RNA survival followed the order: coacervates > PAA only > PDADMA only > no coacervates (plot in Supp. Fig. 6, gels in Supp. Fig. 7-8). We hypothesized that the reduction in RNA hydrolysis in PDADMA-only samples was due to decreased accessibility of RNA’s phosphodiester backbone^41, 42^ upon binding of the polycationic PDADMA to negatively charged RNA (Supp. Fig. 6b). PDADMA’s protective effect reduced with increasing NaCl, consistent with PDADMA – RNA electrostatic interactions. The polycarboxylate, PAA, reduced RNA hydrolysis more than PDADMA, most likely due to its ability to chelate Mg^2+^ (Fig. 3c)^43^. RNA hydrolysis is known to follow an acid-base mechanism^42^, and can be catalyzed by hydrated Mg^2+^ acting as a base (Fig. 3b). Negatively-charged carboxylate ligands on Mg^2+^ could reduce RNA hydrolysis by suppressing deprotonation of the remaining ligated water molecules to generate Mg^2+^–OH^−^ complex^44, 45^. Mg^2+^ chelation by PAA’s carboxylate moieties also decreased at higher salt, presumably due to charge screening by the added NaCl^44–49^ (Supp. Fig. 6a). In sum, although each polyelectrolyte alone offered some degree of protection from hydrolysis, however, neither protected as well as the coacervate samples. Each polyelectrolyte would be expected to interfere with, rather than aid, the other’s protective mechanism, as PDADMA-PAA binding competed with PDADMA-RNA binding or PAA-Mg^2+^ chelation; in other words, their effects should not be additive to explain coacervate effects. Thus, the protective effect observed in coacervate systems cannot be fully explained by the mechanisms available to the individual polyelectrolytes. Finally, the RNA protection effect was largely insensitive to charge imbalance in the coacervate systems over the range tested (1.5- fold excess of either cationic or anionic sidechains; see plot in Supp. Fig. 9 and gels in Supp. 10- 13; similar Mg/Na was observed across charge imbalance samples, see Supp. Fig 14); thus, it would not be necessary to have a 1:1 charge ratio to achieve protection against RNA hydrolysis during wet/dry cycles.

**Figure 6.**
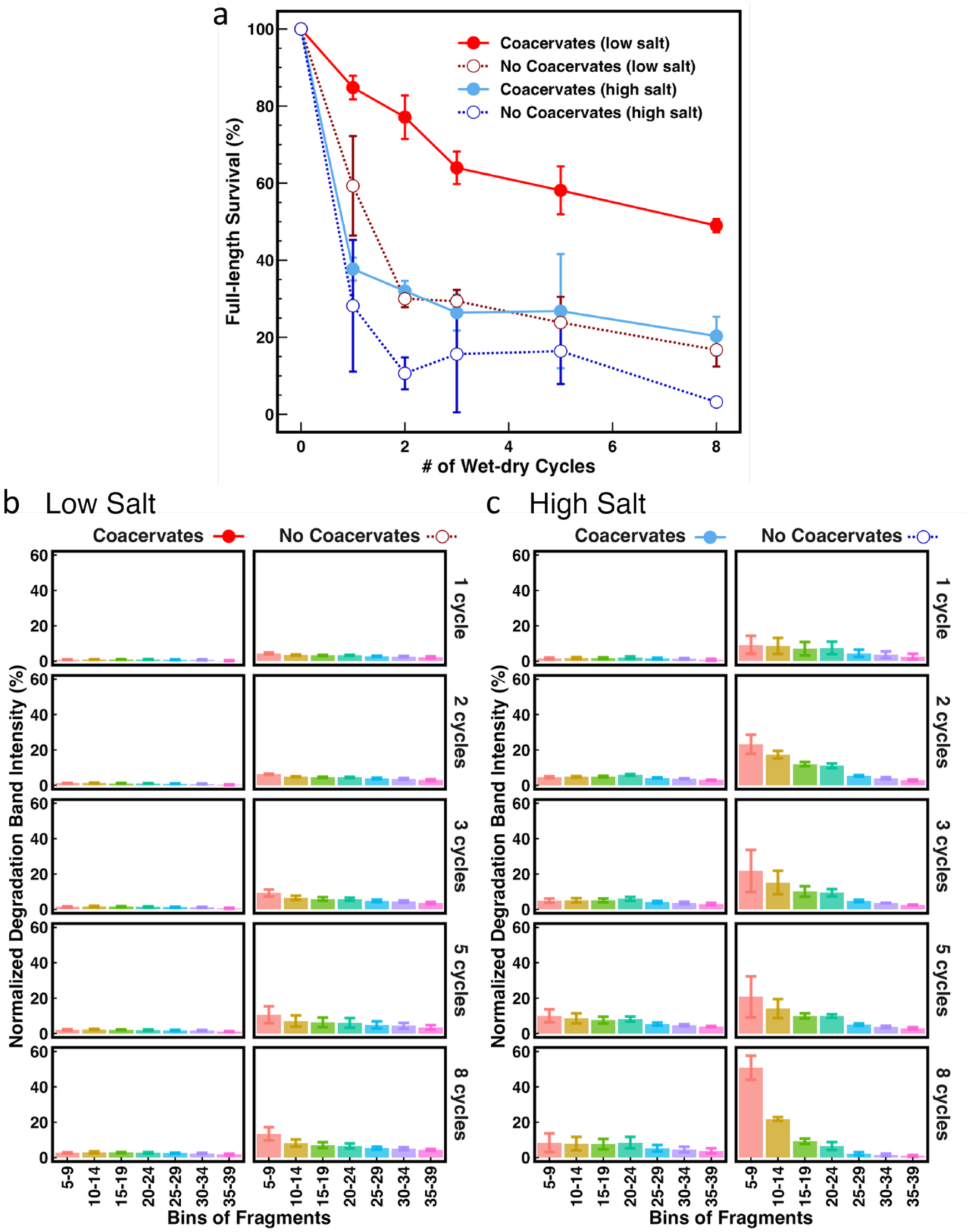
Multiple wet-dry cycles on U45. **a** The full-length U45 survival after 1, 2, 3, 5 and 8 wet-dry cycles within various systems: samples with coacervates under the low salt condition (blue filled square), samples without coacervates under the low salt condition (blue open circle), samples with coacervates under the high salt condition (red filled square), and samples without coacervates under the high salt condition (red open circle). **b** Normalized intensity for degradation bands combined into 5 nucleotide bins under the low salt condition. **c** Normalized intensity for degradation bands combined into 5 nucleotide-bins under the high salt condition. RNA fragments <5 nt are not included because they could not be accurately quantified from the PAGE gels.

**Figure 7.**
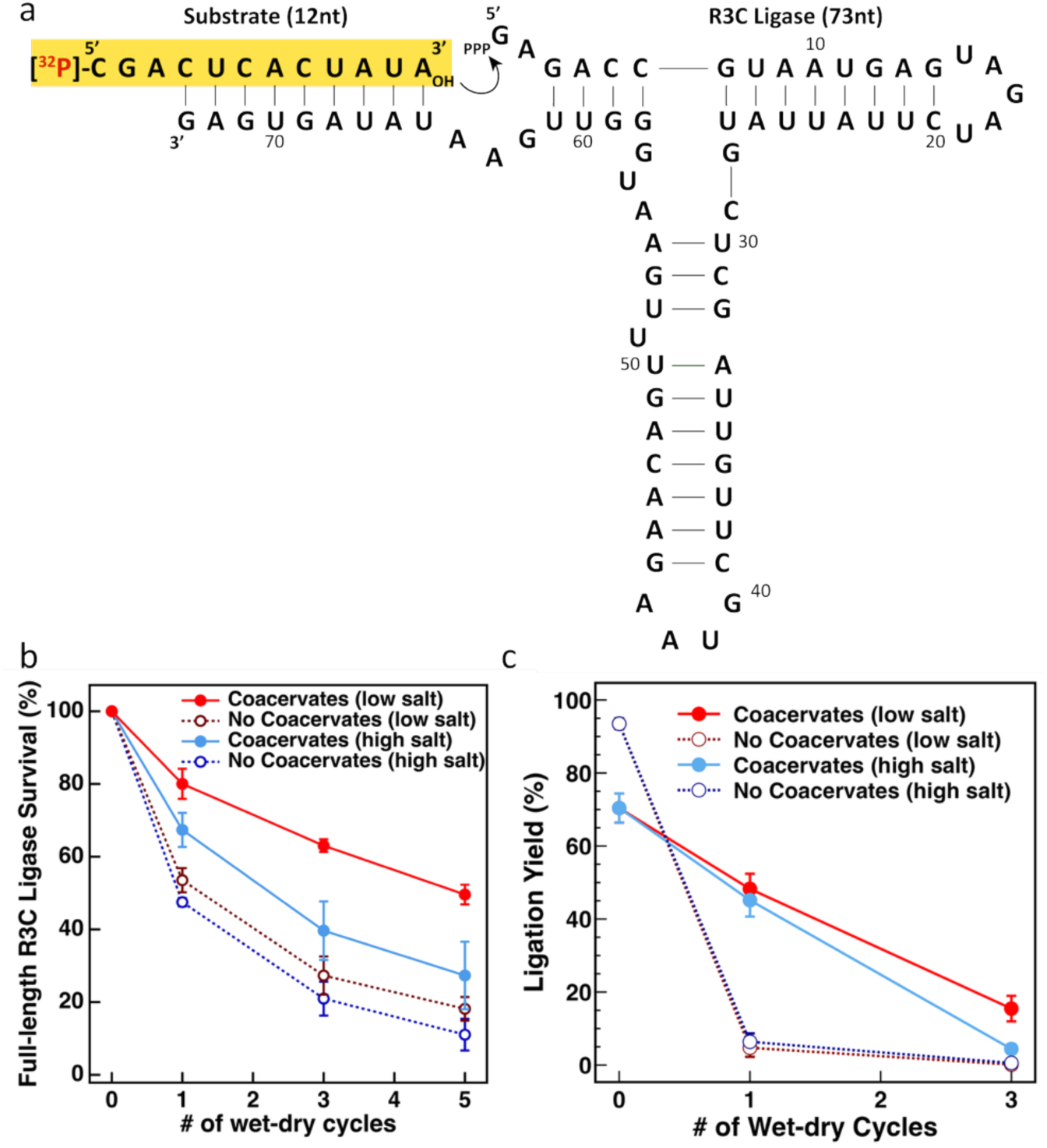
Coacervates protect R3C ligase against hydrolysis throughout wet-dry cycling. **a** Secondary structure of the R3C ligase. It ligates a 12nt substrate (yellow background) to itself with release of pyrophosphate. **b** Full-length survival of 5’-^32^P-labeled R3C ligase after multiple wet-dry cycles at 75 °C. **c** The ligation yield (% of substrate strands that showed increased length after reaction) of after-wet-dry-cycling R3C ligase among different systems. Substrate and ligation buffer (25 mM MgCl_2_ and 40 mM Tris pH = 8.5) were added after samples completed wet-dry cycling. The ligation experiment was conducted at 37 °C for 22 hours using samples straight after wet-dry cycles without additional reannealing steps. The amount of substrate added in the ligation step was the same concentration as R3C ligase before wet-dry cycling. The ligation yield was determined by the % of substrate strand was ligated to R3C ligase. R3C ligase in the ligation reaction had 5’ triphosphate after transcription and no further labeling or treatment.

### Compartmentalization is important for RNA protection against hydrolysis

The coacervate system differs from single-polyelectrolyte or salt controls in its ability to sequester RNA molecules within the polyelectrolyte-rich coacervate droplets, which coexist with the continuous dilute phase but offer a distinct microenvironment. To better understand the role of sequestration within the coacervate phase in protecting RNA, we performed experiments in which the same amount of RNA was added to samples comprised of pure coacervate phase or pure dilute phase (Fig. 4a). To directly compare the effect of heating on RNA molecules located in coacervates vs in dilute phase, there experiments were performed at 75°C for 180 min, but at constant volume (no drying). Under the same salt condition, coacervate phase alone showed similar full-length survival rate as the full two-phase system, which was significantly larger than dilute phase alone or no-polyelectrolyte samples (Fig. 4b, Supp. Fig. 15). These data further support the protective effect of the coacervate environment for RNA. Additionally, the similarly low levels of hydrolysis events in coacervate+dilute and coacervate-only phase samples suggests that these events occur within coacervates rather than requiring RNA excursions into the dilute phase, consistent with the strong sequestration of RNA noted above (>90% of RNA is in coacervates; Fig. 1c, 4c). We also observed the full-length survival percentage was similar for dilute phase alone and no-polyelectrolyte samples, consistent with the low polyelectrolyte concentrations present in dilute phase (Supp. Fig. 15).

Added NaCl concentration was much less important for these constant-volume experiments than for sample drying, with high salt coacervate systems preserving full-length RNA nearly as well as low-salt systems (Fig. 4b; compare to Fig. 3a). This observation underscores the crucial role of coacervate persistence in RNA protection during wet-dry cycling, because without drying the RNA remains compartmentalized within the coacervate phase regardless of the salt concentration and had much more full-length survival (80%-70% in low and high salt under heating constant volume compared to 65%-32% in low and high salt under wet-dry cycling, respectively).

Nonetheless, the low salt condition still showed somewhat better RNA protection than the high salt condition. This can be understood as a consequence of their distinct coacervate phase compositions, with low salt samples having higher polyelectrolyte and lower water concentrations as compared with high salt samples, in accordance with the phase diagram (Fig. 2b, Supp. Fig. 16-18)^24^. Taken together, the drying and constant-volume experiments indicated that compartmentalization within coacervates is responsible for protecting RNA from hydrolysis during wet-dry cycling, and that the main factor distinguishing high and low salt samples is the greater persistence of coacervate phase during drying in the low salt samples, with salt-dependent differences in coacervate phase composition and properties playing a minor role.

### Content and properties of the coacervate phase

Having established that RNA compartmentalization within the coacervate phase was essential, we next sought to understand the organic content, RNA accumulation, and physical properties of the coacervate phase that give rise to its protective effect. Regarding organic content, polyelectrolytes are expected to be greatly enriched within the coacervate phase^53^; with reduced partitioning at elevated ionic strength^17, 29, 54^. We obtained the apparent partition coefficient of polyelectrolytes in our systems using Thermogravimetric Analysis (TGA)^55^. Total organic content, corresponding to PDADMA+PAA, was ∼51× and ∼37× higher in the coacervate than dilute phases at low and high salt, respectively (Supp. Fig. 16-19). MicroRaman supported this trend for our systems, with relative intensities in the C-H stretching region indicating the highest organic content for coacervates prepared in the low salt condition, followed by high salt coacervates, and very little signal for either dilute phase (Figure 1d, Supporting Figure 19b).

Turning to RNA accumulation, as shown qualitatively in Fig. 1e, fluorescently-tagged RNA strongly accumulated within coacervates. We quantified U45 partitioning by scintillation counting using ^32^P 5’-end-labeled RNA, which provides a larger dynamic range than fluorescence and avoids any potential bias of partitioning that can come from a hydrophobic fluorescent tag. Under the low salt condition, the coacervate phase contains 98.5% of the total U45 in just 0.6% of the total volume (Fig. 4c, Supp. Fig. 19; K_U45_ ∼ 1.1× 10^4^, see Methods for *Simulation Calculation*). With increasing NaCl concentration, RNA sequestration within coacervates is less effective, and the percent of total RNA in the coacervate phase drops to 91.6% in 0.5% of the total volume (K_U45_ ∼ 2.4 × 10^3^, see Method for calculation). This difference in RNA partitioning also contributes to the difference in effectiveness in RNA protection between low and high salt conditions, since RNA survival is greater when in the coacervate phase (Fig. 4b-c).

The interior of coacervate droplets is defined not only by the local concentrations of molecules but also by the interaction network between them^29, 56^. We used Fluorescence Recovery After Photobleaching (FRAP) to understand the mobility of U45 in coacervates under the low and high salt conditions^25, 57^. Low salt coacervates showed ∼2-fold slower apparent diffusion of fluorescently tagged U45 in partial-droplet FRAP experiments as compared to high salt coacervates (Supp. Fig. 20), consistent with the higher local polyelectrolyte concentrations and stronger intermolecular binding interactions in low-salt coacervates. Full-droplet recovery times were also markedly slower for low salt as compared to high salt samples, as anticipated from the stronger U45 partitioning and lower availability of RNA outside the droplets (Fi. 4c, Supp. Fig. 21). Another important aspect of coacervate properties is their hydrogen bond network^58^. The hydrogen bonding network of bulk water can be disrupted by dissolved ions (e.g., Na^+^, Cl^−^) and polyelectrolytes^50, 59, 60^. Systems with greater hydrogen bonding tend to have higher 3200 cm^-1^ peak intensity (low frequency water shoulder)^59, 61^. Our microRaman data showed less tight bonded water not only for the high salt vs low salt samples, as expected, but also in the coacervate interior as compared to its surrounding dilute phase under the same salt conditions (Fig. 4d, Supp. Fig. 18b). The relative reduction in this ∼3200 cm^-1^ feature for coacervates as compared to their dilute phases is consistent with the higher local polyelectrolyte concentration of the coacervate phase, where increased interaction between water and electrolytes disrupts the hydrogen bonding network and water activity is reduced^50, 59, 60, 62^.

### Mg^2+^ accumulation and availability within coacervates

Next, we turned to Mg^2+^ concentration because it plays a critical role in RNA degradation. ICP-AES revealed 90-100 mM Mg^2+^ within the coacervate phase, much higher than the 2 mM overall Mg^2+^ present in these samples. Mg^2+^ partition coefficients (K_Mg_) were relatively insensitive to added NaCl concentrations, with lowest and highest values of ∼60 and ∼75 at 0 and 50 mM added NaCl, respectively (Fig. 5a, Supp. Fig. 22). Accumulation of Mg^2+^ is driven by binding to PAA’s carboxylate moieties, which are also greatly concentrated within the coacervate phase^26^. It is striking that the coacervate phase is strongly protective against RNA hydrolysis despite harboring much higher local concentrations of Mg^2+^, which is a potent catalyst for RNA hydrolysis^6, 63^, indicating that this Mg^2+^ is less available to interact with RNA, and/or that the pKa for Mg^2+^-bound H_2_O has been shifted by its coordination to PAA’s carboxylates^44, 45^. Na^+^ concentrations in the coacervate and dilute phases were similar, with K_Na_ ranging from ∼0.6 to ∼1.1 between low and high salt conditions (Supp. Fig. 22 b-c). Increased relative amounts of Na^+^ vs Mg^2+^ in the coacervate phase with increasing added salt (Fig. 5a) effectively weakens the chelation of Mg^2+^ by coacervate carboxylate moieties, potentially contributing to the less effective protection against RNA hydrolysis at high salt.

We further explored the importance of Mg^2+^ chelation within coacervates by adding a strong chelator – ethylenediaminetetraacetic acid (EDTA) and by swapping PAA for an equal charge concentration of propionic acid (PA) (Supp. Fig. 23-25), which has only one carboxylate moiety. Adding EDTA or omitting Mg^2+^ improved full-length RNA survival similarly under both salt conditions (Fig. 5b). EDTA can replace all the H_2_O from Mg^2+^ hydration shell, disabling hydroxyl production to suppress RNA hydrolysis^64^. PAA is unable to achieve this level of Mg^2+^ sequestration; however, it does greatly outperform the small molecule chelator, PA, at both salt concentrations and in the presence or absence of PDADMA (Figure 5c, Supp. Fig. 25)^65^. We note that unlike PAA, neither EDTA nor PA formed coacervates with PDADMA under our experimental conditions, hence they were unable to provide compartments for RNA.

### Initial polyelectrolyte concentrations control RNA protection during dehydration: Less is more

As water evaporates, the composition of the coacervate system changes according to the phase diagram (Fig. 2b). Somewhat counterintuitively, as the system dehydrates, the coacervate phase becomes less concentrated in polymers, and more similar in composition to the supernatant phase before it ultimately disappears^24^. Consequently, the partitioning of solutes such as Na^+^, Mg^2+^ and RNA does not remain constant during drying. We therefore prepared matched-volume “drying- mimic” samples with compositions matching samples evaporated to 70% of their initial volume, which was chosen as a substantial volume change that was still well within the two-phase region for both low- and high-salt drying trajectories (Fig. 2b). For low salt samples, > 97% of the total RNA remained encapsulated within the coacervate phase in the 70% volume drying-mimic samples (Fig. 4c, Supp. Fig. 26). In contrast, the same volume reduction resulted in a decrease in encapsulated RNA from ∼92% to ∼79% for high salt samples (Fig. 4c). Thus, even before the high salt coacervates dissolve at ∼50% volume, RNA is less well sequestered.

We also measured the local concentrations for Mg^2+^ and Na^+^ in the 70% volume drying mimic samples. We found that Mg^2+^ partitioning was markedly reduced in both high and low salt drying mimics, more steeply so in the high salt samples (Supp Fig. 27a). The [Mg^2+^]/[Na^+^] ratio in coacervates was also impacted. For the low salt condition, [Mg^2+^]/[Na^+^] decreased from ∼6.5 to ∼4.0 (Supp. Fig. 27b), however it still remained substantially higher for the low salt than the high salt condition, for which this ratio was <1.

Less effective sequestration of RNA and an influx of Na^+^ suggested that the coacervate protection effect likely becomes weaker as the drying process progresses. This result made us hypothesize that for different initial coacervate samples located on the same drying trajectory, initial compositions closer to the origin of the phase diagram would offer superior protection of RNA against hydrolysis (Fig. 2b), benefitting not only from greater coacervate persistence during drying but also from better coacervate sequestration capabilities while present. To test this idea, we prepared samples of coacervate systems with different dilution factors, such that their initial points fell on the same drying trajectory (Supp. Fig. 28a). Indeed, samples whose initial compositions were closer to the origin of the phase diagram showed higher full-length U45 RNA survival under both salt conditions tested (Supp. Fig. 28-30). Despite the charge concentration difference, the low salt condition always performed better in protecting full-length U45. These data further supported our understanding of the mechanisms for coacervate protection against RNA hydrolysis and provided guidance on how to best optimize the protection for a given polyelectrolyte combination. Since complex coacervate phase behavior follows the same general trends across a wide variety of polyelectrolyte chemistries^25, 56^, we expect these findings to apply broadly, well beyond the PDADMA/PAA system studied here. With respect to prebiotic compartmentalization, these findings suggest that to protect RNA on the early Earth under wet-dry cycling, there was no need to have high concentrations of polyelectrolytes, as long as the coacervate compartments were present. Indeed, low polyelectrolyte concentrations could benefit RNA more in protection.

### RNA hydrolysis under multiple wet-dry cycles

Performing multiple wet-dry cycles provides additional opportunities for hydrolysis and loss of full-length RNA (Fig. 6a, Supp. Fig. 31-32). General trends from single wet-dry cycle persist: coacervate-containing samples retain more full-length RNA than no-coacervate samples, and low-salt coacervate samples provide the greatest protection with nearly 50% of full-length U45 RNA remaining after 8 cycles, as compared with only ∼20% for the coacervate-free low salt sample. High salt samples only preserved ∼15% of full-length RNA with coacervates, and a mere 3% without coacervates.

Full length RNA survival is only part of the story. We noticed that only coacervate samples under the low salt condition showed a linear relationship between the full-length survival rate and number of wet-dry cycles. For other conditions, it seemed that the full-length survival rate declined substantially after one or two cycles albeit less so after three cycles, when less than ∼35% full-length RNA survived.

During repeated wet-dry cycles the likelihood of multiple cleavage events on a single strand of RNA increased. Eventually the population of RNA fragments shifted to shorter and shorter lengths; this is most clear for the high salt condition without coacervates (Fig.6b-c, Supp. Fig. 31-36). The accumulation of short RNA fragments is directly related to the increasing probability of phosphodiester bond cleavage.

We were curious whether the probability of cleaving any individual phosphodiester bond depended on RNA fragment length. Although polyU is considered to be a single stranded and random coil under room temperature^40^, suggesting all bonds will have the same cleavage susceptibility, we were curious whether stronger sequestration of longer RNAs would result in their preferential protection. Comparing U15 and U45, we observed higher survival for the U15 consistent with its smaller number of potential cleavage sites and not preferential protection of longer RNA (Supp. Fig. 37-39). To better understand cleavage probabilities at each site, we simulated RNA degradation fragment distributions assuming each bond has the same cleavage probability regardless of its location within any RNA fragment (details of the simulation can be found in the Supporting Information.) Per bond cleavage probabilities (X) able to produce RNA fragmentation patterns matching the experimental data were identified for all conditions, supporting the assumption of unbiased hydrolysis across all phosphodiester bonds. X values for a single wet-dry cycle in the absence of coacervates were found to be 0.0069 ± 0.0003 and 0.016 ± 0.005 at low and high salt, respectively. With coacervates, these values dropped by more than 4- fold, to 0.0016 ± 0.0001 at low salt and by more than 6-fold, 0.0025 ± 0.0004, at high salt.

(Supp. Fig. 40-42). Trends in bond cleavage probability followed those of apparent full-length survival (Fig. 6a, Supp. Fig. 40-42). Using these per-bond cleavage probabilities, we then simulated a full degradation fragment probability distribution that included fragments below 5 nts, which could not be directly quantified from experimental data (Supp. Fig. 40, 42). Despite differences in the full-length survival of U45 and U15 (Supp. Fig. 37), their per-bond cleavage probabilities were similar (Supp. Fig. 44).

These data show that coacervates not only preserve full-length RNA strands through wet-dry cycling but also help prevent formation of ever-shorter RNA fragments by multiple cleavages. All phosphodiester bonds showed equal hydrolysis probability regardless of their location within the larger U45 (or U15) strand. Although longer RNAs are sequestered more fully, it was possible to simulate fragmentation patterns matching our experimental data without including a length bias. Therefore, any differences due to weaker partitioning of shorter RNAs in our system were small, probably because even the relatively short U15 RNA was well-sequestered in these PDADMA/PAA coacervates^24, 27^. In the context of prebiotic wet-dry cycling, coacervates, by maintaining a greater population of longer RNAs, could help support productive oligomerization and ribozyme activity.

### Coacervate protection of a prebiotically plausible ligase ribozyme

Could the protection from hydrolysis afforded by encapsulation within coacervates have aided prebiotic catalysis by ribozymes? To test this, we chose a ribozyme ligase because ligases avoid the confounding problem of sorting out ribozyme cleavage from background cleavage. The R3C ribozyme ligase^66^ (Fig. 7a) is one of the shortest ligases (73nt) that can perform self-ligation, a potentially important primitive ribozyme activity, and can tolerate multiple mutations while maintaining ligation function^2, 66, 67^. Although the R3C ligase is structured at room temperature (Fig. 7a), we expect it to be unfolded during drying at 75°C (Tm ∼50°C predicted by RNAfold). Survival of full-length R3C ligase through multiple wet/dry cycles followed the same trends as U45, with coacervates more than doubling the amount of full-length ribozyme remaining after 5 wet-dry cycles, and low-salt coacervate systems offering maximal protection (Fig. 7b, Supp. Fig. 45-46). Although the presence of coacervate components reduced R3C ligation yields from ∼95% to ∼70% when no wet-dry cycling was performed, coacervate-containing ligase samples greatly outperformed no-coacervate samples after wet-dry cycling (Fig. 7c, Supp. Fig. 47-48). Specifically, a single wet-dry cycle essentially inactivated the ligase in no-coacervate samples, while yields were ∼50% for both low and high salt coacervate-containing samples.

Two aspects of these data stand out: First, for no-coacervate samples, the loss of ribozyme activity was much greater than the loss of full-length R3C. Second, ribozyme function was surprisingly insensitive to the low vs high salt condition during wet-dry cycling, despite the greater preservation of full-length R3C ligase in low-salt samples (Fig. 7b). We hypothesize that these observations could be due to coacervate-based protection from additional types of damage beyond backbone cleavage that could also occur during wet-dry cycling at 75°C. Coacervates may protect against loss of the ribozyme’s terminal triphosphate, which is required for the ligation reaction, and is expected to undergo hydrolysis more readily than RNA’s phosphodiester bonds.

## Conclusion

We have demonstrated that complex coacervates formed from oppositely charged polyelectrolytes protect RNA from Mg^2+^-catalyzed hydrolysis. By compartmentalizing within coacervates, a much greater fraction of full-length RNA can survive several wet-dry cycles, and larger fragments persist through more wet-dry cycles than in the absence of coacervates. These findings support the potential of coacervates as prebiotic compartments for protection of RNA or similar hydrolysis-prone protobiomolecules in the emergence of life. Coacervates have previously been shown to support RNA folding^28^. Thus, compared with stronger Mg^2+^ chelators such as EDTA, which can also protect RNA from hydrolysis, carboxylate-rich coacervates offer RNA sequestration within a distinct microenvironment that not only provides hydrolysis protection, as demonstrated here, but also can support the adoption of Mg^2+^-dependent functional RNA secondary and tertiary structures. In this way, coacervate-based compartments can provide a hospitable environment for ribozymes while offering protection from hydrolysis during wet-dry cycling.

Beyond its importance for the emergence of life, extending RNA’s structural and functional integrity is also relevant for biotechnology, for example in constructing more robust artificial cells and improving the stability of RNA reagents and therapeutics. Our findings suggest the possibility that compartmentalization of RNA within coacervates, already of interest for packaging and delivery^68^, could additionally be employed to protect encapsulated RNAs during transport and storage^69^.

## Acknowledgements

This project was supported by the NASA Exobiology program Grant No. 80NSSC22K0553. The co-authors acknowledge the Pennsylvania State University Materials Characterization Lab Core Facility, Materials Research Institute, University Park, PA (RRID:SCR_012386) for use of microRaman and Dr. Maxwell Wetherington for assistance with microRaman data collection and analysis. We thank Kobie Kirven for checking the simulation calculation.

## Supporting Information

Supplementary figures, tables and methods will be made available upon peer review.

